# Leukocyte adhesion is governed by endolysosomal two pore channel 2 (TPC2)

**DOI:** 10.1101/2021.09.28.462104

**Authors:** Jonas Goretzko, Nicole Heitzig, Katharina Thomas, Einar Kleinhans Krogsaeter, Johannes Naß, Anna Lívia Linard Matos, Tristan Wegner, Sebastian Schloer, Volker Gerke, Jan Rossaint, Frank Glorius, Franz Bracher, Christian Grimm, Ursula Rescher

**Affiliations:** Research Group Regulatory Mechanisms of Inflammation, Institute of Medical Biochemistry, Center for Molecular Biology of Inflammation, University of Muenster, von-Esmarch-Str. 56, 48149 Muenster, Germany; Faculty of Chemistry / Biochemistry III, University of Bielefeld, Universitaetsstr. 25, 33615 Bielefeld, Germany; Department of Anesthesiology, Intensive Care and Pain Medicine, University Hospital Muenster, Albert Schweitzer Campus 1, A1, 48149 Muenster, Germany; Walther-Straub-Institute for Pharmacology and Toxicology, Ludwig-Maximilians-University, Nussbaumstr. 26, 80336 Munich, Germany; Institute of Medical Biochemistry, Center for Molecular Biology of Inflammation, University of Muenster, von-Esmarch-Str. 56, 48149 Muenster, Germany; Institute of Organic Chemistry, University of Muenster, Corrensstr. 40, 48149 Muenster, Germany; Department of Pharmacy, Center for Drug Research, Ludwig-Maximilians-University, Butenandtstr. 5-13, 81377 Munich, Germany

**Keywords:** CD63, endolysosome, leukocyte recruitment, TPC2, Weibel-Palade body

## Abstract

In response to pro-inflammatory challenges including pathogenic attack and tissue damage, the endothelial cell surface is rearranged to present leukocyte-engaging cell surface receptors. The initial contact needed for leukocyte tethering and rolling is mediated via adhesion demand-driven exocytosis of Weibel-Palade bodies (WPB) that contain the leukocyte receptor P-selectin together with the stabilizing co-factor CD63. We found that diminished expression of the endolysosomal non-selective cation channel TPC2 or inhibition of TPC2-mediated Ca^2+^-release via trans-Ned 19 led to reduced endolysosomal Ca^2+^ efflux, and blocked transfer of CD63 from late endosomes/lysosomes (LEL) to WPB, and a concomitant loss of P-selectin on the endothelial cell surface. Accordingly, P-selectin-mediated leukocyte recruitment to trans-Ned 19-treated HUVEC under flow was significantly reduced without disturbing VWF exocytosis. Our findings establish the endolysosome-related TPC2 Ca^2+^ channel as a key element in the maintenance of proper endothelial functions and a potential pharmacological target in the control of inflammatory leukocyte recruitment.

## Introduction

While leukocytes are carried along with the flow of blood under normal conditions, the adhesion of leukocytes to the endothelium is a hallmark of the inflammatory response and initiates leukocyte migration across the endothelial barrier into the tissue (Phillipson & Kubes, 2011; Rossaint & Zarbock, 2013). Upon inflammatory activation, endothelial cells increase the cell surface exposure of selectins, causing leukocyte tethering and rolling along the vessel wall, which is a prerequisite for leukocyte activation. Next, leukocyte integrins ensure firm binding to the endothelial surface, leading to leukocyte arrest and subsequent transendothelial migration. Thus, effective regulation of initial leukocyte attraction to the endothelial surface demands the precise regulation of supramolecular leukocyte docking structures on the endothelial cell surface. The adhesion molecule P-selectin, together with the tetraspanin CD63 as co-factor, serves as the leukocyte receptor in early leukocyte recruitment (Poeter *et al*, 2014; Doyle *et al*, 2011; McEver, 2015). P-selectin and CD63 are stored in unique endothelial secretory organelles, the Weibel Palade bodies (WPB) (Harrison-Lavoie *et al*, 2006). Once the endothelium is stimulated by inflammatory mediators, these lysosome-related granules fuse with the plasma membrane and increase CD63/P-selectin surface expression (Holthenrich *et al*, 2019). Thus, demand-driven exocytosis of these adhesion molecules from intracellular storage compartments ensures the immediate early leukocyte capture.

According to their task as endothelial emergency kits, WPB contain a range of factors that are needed for proper endothelial function. The marker cargo is von Willebrand factor (VWF), a large multimeric glycoprotein that mediates platelet adhesion and fibrin formation at sites of vascular injury (Schillemans *et al*, 2019). After biosynthesis, VWF multimers are assembled in the trans-Golgi network, from where the densely-packed VWF tubules bud to form immature WPB. Although their effective delivery to WPB at appropriate levels is essential, P-selectin and CD63 are sorted onto WPB at different stages of WPB genesis. While newly synthesized P-selectin is included already during WPB formation due to direct binding to VWF, CD63 is trafficked from endolysosomes to the mature WPB (McCormack *et al*, 2017; Harrison-Lavoie *et al*, 2006; Poeter *et al*, 2014). Altogether, it is clear that cellular sorting systems operate to control CD63 levels in WPB and ultimately regulate endothelial responses

A well-established model system for inflammatory-driven WPB exocytosis are human primary endothelial cells derived from the umbilical vein (HUVEC). In these cells, WPB are readily detectable through their characteristic elongated shape and VWF staining, and agonist-driven WPB exocytosis can be followed. Using this model system, we have previously shown that proper sorting of CD63 from endolysosomes onto WPB is critical for proper leukocyte attraction to the activated endothelium (Poeter *et al*, 2014). In search for factors that control LEL to WBP transport route, we here concentrate on the two pore channel (TPC) members TPC1 and TPC2, as they localize to endolysosomes and have been associated with endosomal functions (Marchant & Patel, 2015; Grimm *et al*, 2017). We found that diminished TPC2 expression or inhibition of TPC2-mediated Ca^2+^-release by trans-Ned 19 led to reduced endolysosomal Ca^2+^ efflux, a blocked transfer of CD63 from LEL to WPB, and a concomitant loss of P-selectin on the endothelial cell surface. Accordingly, leukocyte recruitment to trans-Ned 19-treated HUVEC was significantly reduced without significantly disturbing VWF exocytosis. These findings identify the endolysosomal channel TPC2 as a key operational element in the establishment of endothelial leukocyte attraction.

## Results

### TPC1/2 is required for endolysosomal maintenance

TPC are located on membranes of acidic endolysosomes and both of the two human TPC homologs, TPC1 and TPC2, are robustly expressed in HUVEC (Figure 1A). Whereas TPC2 is reported to predominantly localize to endolysosomes, TPC1 is associated with less acidic early endosomes (Morgan *et al*, 2011; Zhu *et al*, 2010). We, therefore, explored whether TPC1 and TPC2 have overlapping or distinct endosomal localization profiles in HUVEC. We transfected HUVEC with expression plasmids encoding for TPC1 and TPC2. As shown in Fig. 1B, TPC1-positive endosomes were much more scattered throughout the cell, similar to internalized transferrin, whereas TPC2 localized almost exclusively to CD63-positive perinuclear endosomes. Interestingly, neither channel was found on WPB, the specialized lysosome-related organelles found in HUVEC (Supp. Fig. 1)

**Figure 1.**
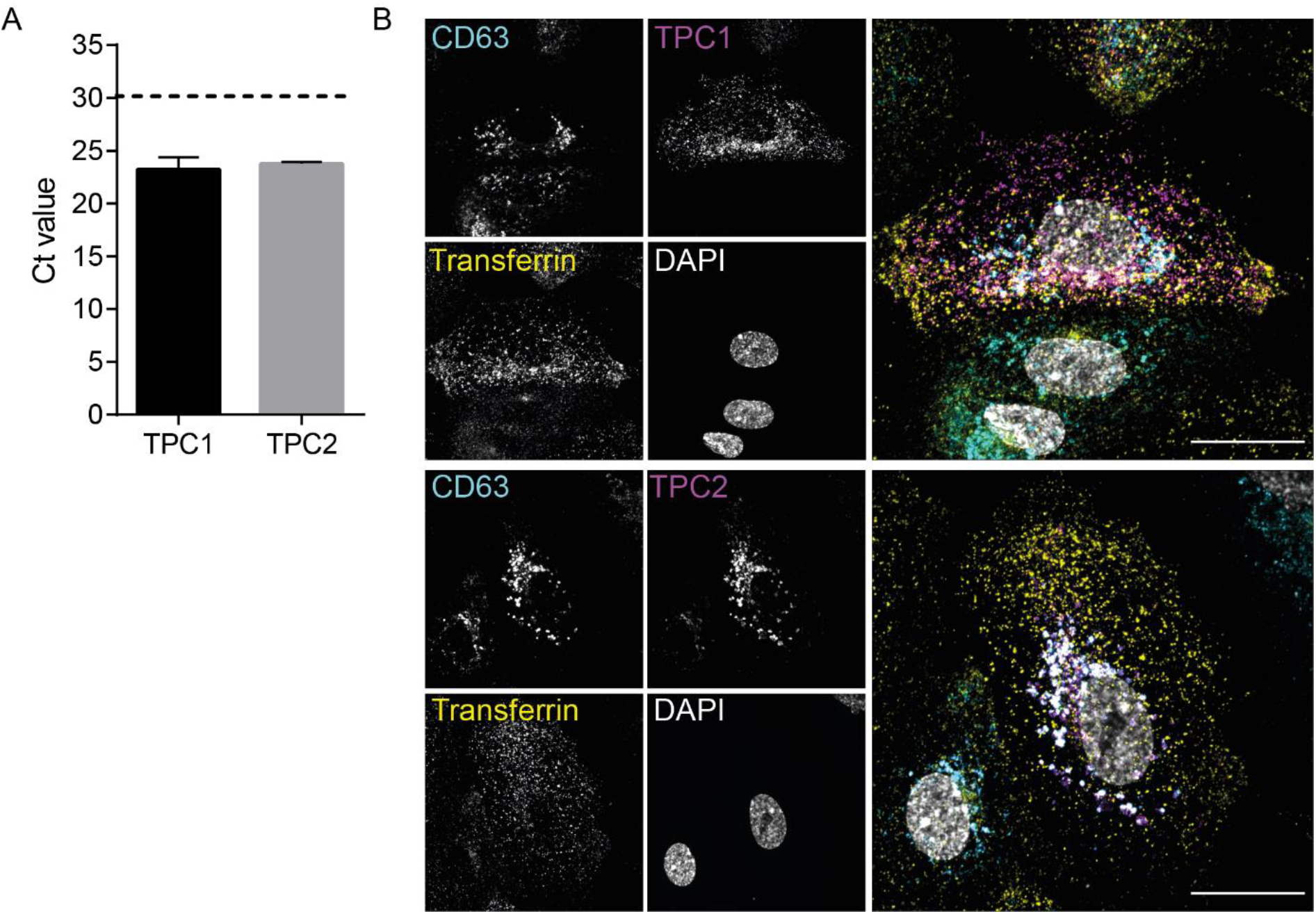
Expression and localization of endosomal two pore channels in HUVEC. (A) The mRNA expression levels of the cation-selective ion channel TPC 1 and 2 were analyzed by qPCR. The threshold cycle value (Ct) considered relevant for protein expression was set to 30 (dotted line). Data represent means ± SEM from four independent experiments. (B) HUVEC transfected with HsTPC1:mRFP and HsTPC2(L564P):mCherry constructs (magenta), subjected to transferrin-AF488 (30 min uptake, yellow) and for stained late endosomal marker CD63 (blue) and nucleus (grey). Images were taken with an LSM 800 airyscan microscope equipped with a 63x oil-objective (Zeiss). Scale bar: 20 µm.

To explore the impact of TPC2 function on endolysosomal integrity, we next assessed the endolysosome morphology in HUVEC downregulated for these channels via siRNA. Because the knockdown of TPC1 or TPC2 has been reported to amplify the expression of the other remaining channel (Jahidin *et al*, 2016), both TPC1 and TPC2 were simultaneously downregulated (indicated as siTPC). RNAi treatment with sequence-specific siRNA duplexes reduced mRNA expression of TPC1 to 25%, and TPC2 to 50%. The successful knockdowns were also confirmed at the protein level (Suppl. Fig. S2). To assess the impact of these channels on endolysosomal morphology, we next visualized endolysosomes immunostained for the specific markers Lamp2 and CD63 via confocal microscopy. In TPC1/2-silenced cells, endolysosomes were enlarged and displayed accumulated CD63 levels (Fig. 2A). To confirm these effects in a quantitative manner, we employed maximum projection to extract and combine the high-intensity structures of both markers from the z-stack volumetric data into single bidimensional images. Matching the images, TPC depletion correlated with an increase in the total LAMP2-positive area per cell (Fig. 2B), and an increase in the integrated CD63 density detected in the Lamp2-positve areas (Fig. 2C), pointing toward an aberrant CD63 retention and indicating that TPC has a direct role in the control of the endolysosomal maintenance.

**Figure 2:**
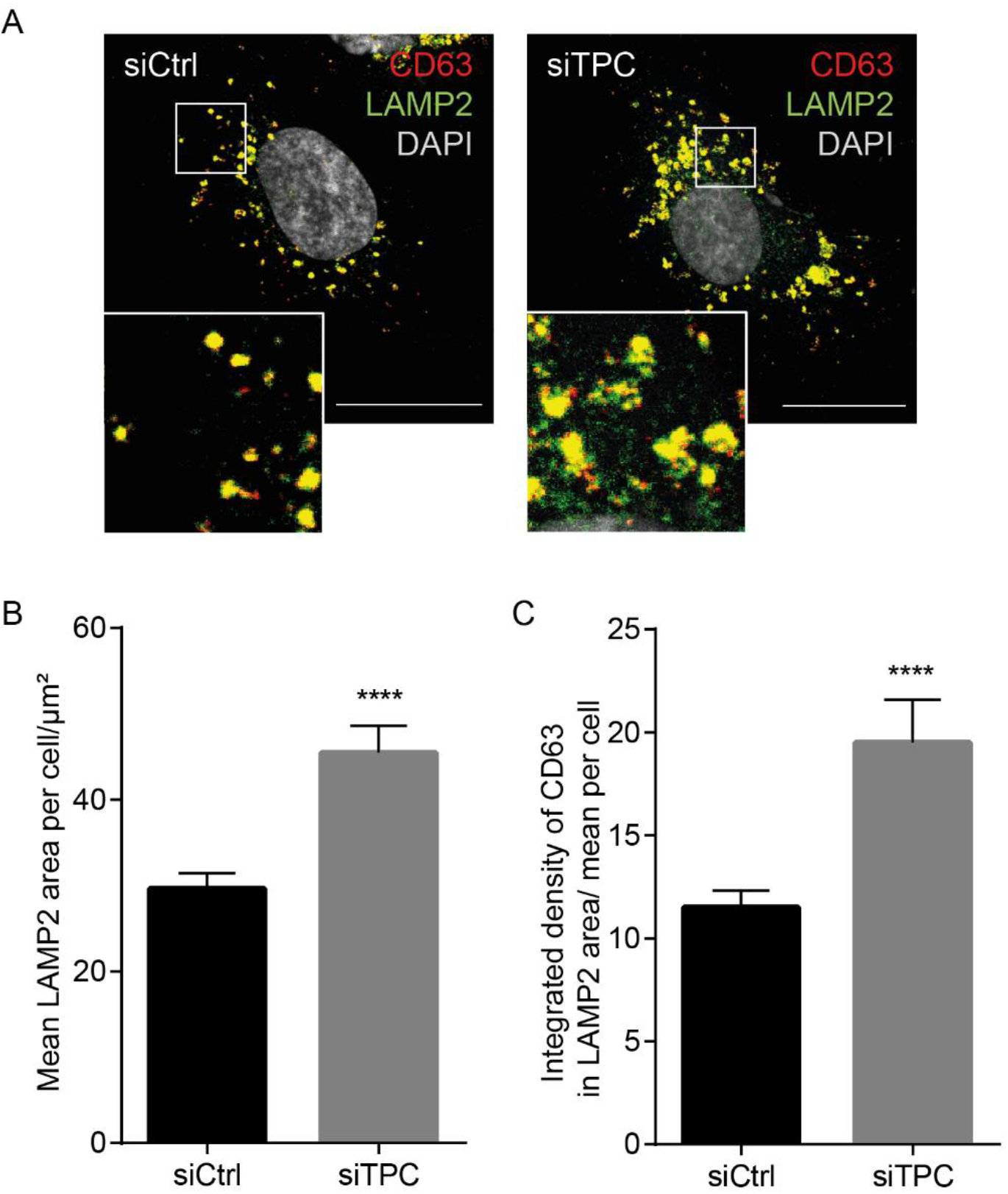
TPC1/2 deficiency alters endolysosomal morphology. (A) Confocal images of HUVEC transfected with either non-targeting siRNA (siCtrol) or siRNA specific to TPC1/2 and immunostained for the endolysosomal marker proteins CD63 (red) and Lamp2 (green). Nuclei were visualized with DAPI (grey). Boxed areas indicate the regions magnified in the insets. Scale bars, 20 µm. (B) Size analysis of Lamp2 positive particles derived from maximum projections of z-stacks of individual cells, (C) Quantification of the CD63 density within LAMP2-positive areas. Data are presented as mean ± SEM of at least 45 cells from three independent experiments, statistical analysis was performed using a two-tailed unpaired Student’s *t*-test (****P < 0 .0001).

### TPC2 function can be pharmacology addressed in HUVEC

Based on the specific endolysosomal localization, we focused on TPC2. To verify that TPC2 on HUVEC endolysosomes was indeed functionally active, we intended to record TPC2 activation in live cells. Toward this goal, we fused the constitutively fluorescent red fluorescent protein mApple to the TPC2-based genetically encoded Ca^2+^ indicator TPC2:GCaMP6 (Gerndt *et al*, 2020), thus generating TPC2-wt-GCaMP6-mApple, a ratiometric indicator to monitor TPC2-mediated Ca^2+^ alterations in real-time, with the additional advantage of seeing TPC2 localization in resting cells. As a specific control we generated a corresponding non-active pore-dead TPC2 mutant construct, TPC2-pore-dead-GCaMP6-mApple. To specifically address the TPC2 activity state in vivo, we employed the recently developed pharmacologic TPC2 activator TPC2-A1-N (Gerndt *et al*, 2020). Imaging the mApple-transmitted signals confirmed that both the active and a pore-dead TPC2 protein resided on late endosomes. Ratiometric tracing revealed that the basal TPC2 activity in resting cells was very low but could be specifically activated via stimulation with the TPC2-A1-N activator which caused a rapid peak and a prolonged activity above baseline levels (Fig. 3A). Stimulation of cells transfected to express the pore-dead TPC2 did not result in any detectable changes in Ca^2+^ levels, thus confirming that the TPC2-A1-N-mediated emission shift was indeed a function of TPC2-mediated increase in local Ca^2+^ concentration (Fig. 3B-C).

**Fig. 3.**
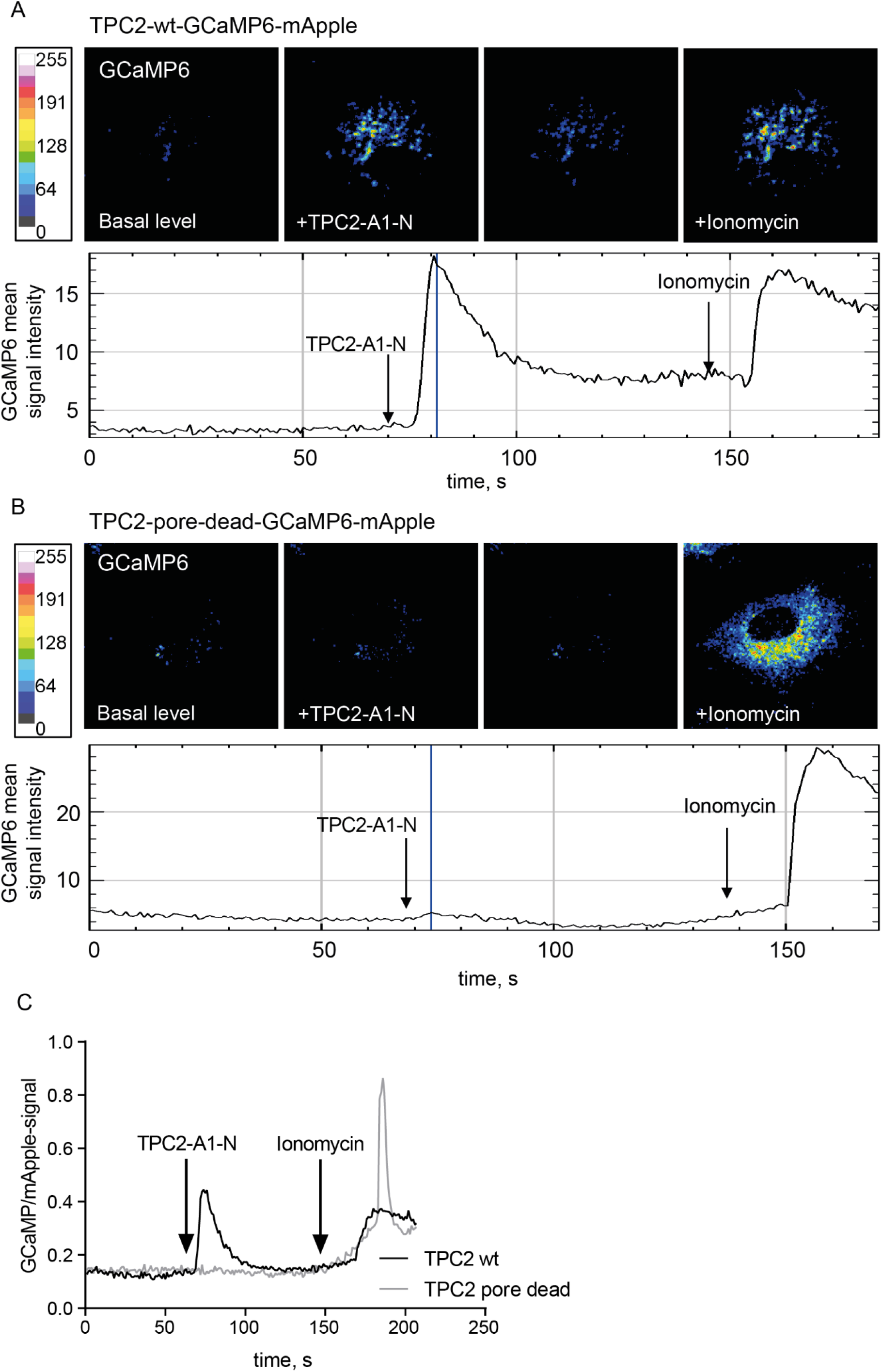
TPC2-specific activation with TPC2-A1-N as determined via a TPC2-based ratiometric Ca^2+^ indicator. Representative confocal images of wild-type (A) and pore-dead (B) TPC2-GCaMP-mApple constructs expressed in HUVEC. GCaMP fluorescence signal is elicited upon stimulation with the TPC2 activator TPC2-A1-N and signal intensity is depicted as a heatmap. (C) Comparison of representative time-resolved fluorescence ratios in TPC2-A1-N stimulated cells expressing either the active TPC2 (black trace) or the pore-dead TPC2 (grey trace). Note that ionomycin-induced unspecific elevation of cytosolic Ca^2+^ levels was still observable in both cases.

We next traced the endolysosomal Ca^2+^ release in the presence of the TPC2 inhibitor trans-Ned19. Because endolysosomal Ca^2+^ release is local and rapidly sequestered, we added bafilomycin A1 (BafA1), an inhibitor of the V-ATPase, to prevent endolysosomal Ca^2+^ reuptake, thus amplifying the changes in cytosolic Ca^2+^ levels that arose from endolysosomal Ca^2+^ release (Lloyd-Evans *et al*, 2008; Davis *et al*, 2012; Morgan *et al*, 2020; Yang *et al*, 2019; Morgan & Galione, 2021). As expected, BafA1 addition increased cytosolic Ca^2+^ levels in control cells. In trans-Ned 19-treated cells, the increase was significantly smaller, strongly suggesting that the TPC2 contribution was inhibited (Fig. 4A-B).

**Figure 4.**
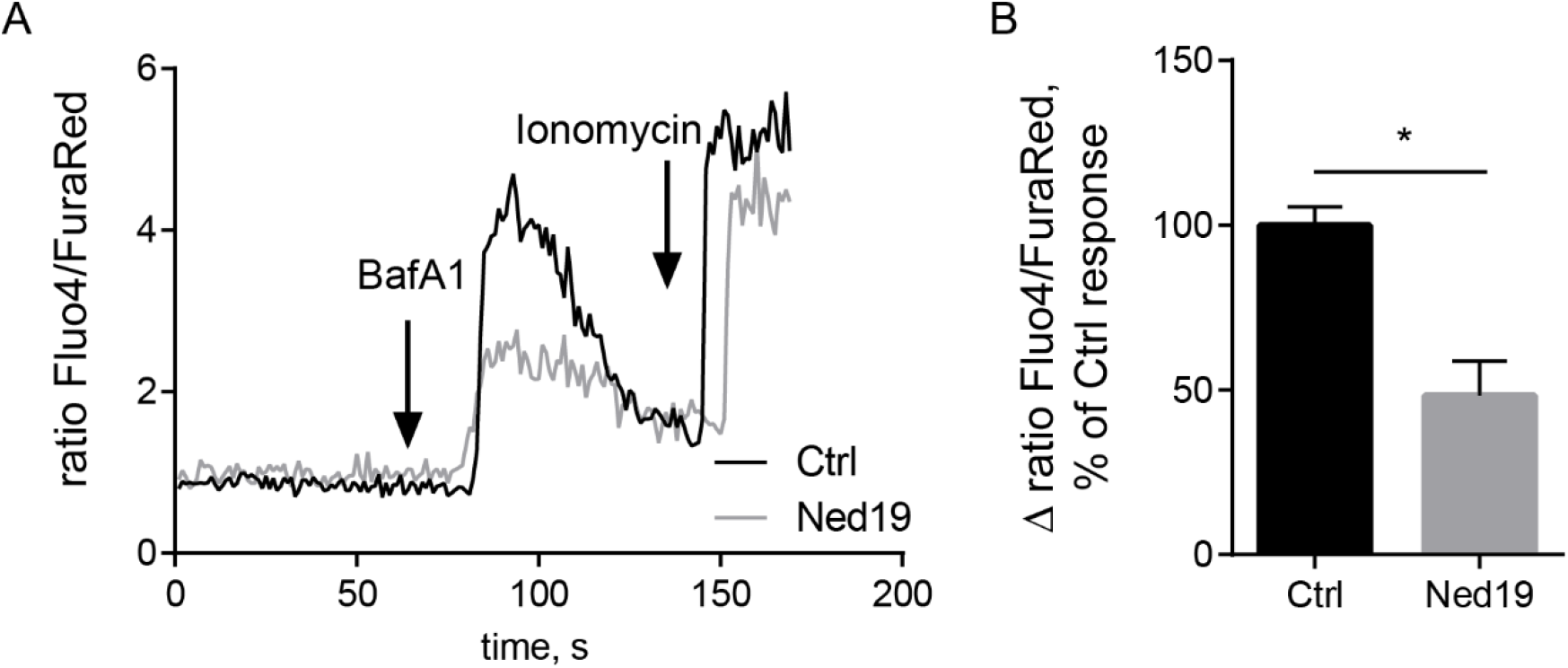
TPC2 inhibition impairs endolysosomal Ca^2+^ release. Ratiometric in vivo Ca^2+^ measurements showing a decreased Ca^2+^ release from endolysosomal stores of cells treated with the TPC2 inhibitor trans-Ned 19 (10 µM). Blocking lysosomal Ca^2+^ reuptake with the VATPase inhibitor bafilomycin A1 (250 nM) was used to amplify Ca^2+^ signals. (A) Representative ratiometric time course measurement of endolysosomal Ca^2+^ release upon bafilomycin A1 stimulation of control cells and trans-Ned19 treated cells. (B) Quantification of Ca^2+^ release of 24 cells per condition from four independent experiments. Ca^2+^ response is depicted in relation to the control. Two-tailed Student’s t-test (*p<0,05) was performed on the raw data.

### TPC2 is required for endolysosomal cholesterol balance

TPC2 deficiency in mice has been shown to cause defective endolysosomal cholesterol handling (Grimm *et al*, 2014). Because we found endolysosomal cholesterol balance to govern fundamental endothelial cellular functions (Heitzig *et al*, 2017, 2018; Poeter *et al*, 2014) we, therefore, assessed whether the profoundly changed endolysosomal morphology in TPC2-targeted HUVEC was accompanied by an accumulation of cholesterol in the enlarged endolysosomes. Thus, we treated cells with either TPC1-A1-N or trans-Ned 19 and also included U18666A (U18), a widely used direct inhibitor of the endolysosomal cholesterol transporter NPC1, as an internal reference for maximum system response. To visualize cholesterol, we used the fluorescent polyene macrolide antibiotic and antifungal filipin that binds to membrane cholesterol and is commonly used to track cellular cholesterol (Gimpl & Gehrig-Burger, 2007) (Fig 5A). Confocal analysis and quantitative colocalization analysis of z-stacks of individual cells (Fig. 5C) revealed that the cholesterol content in CD63-positive endosomes was elevated in trans-Ned 19 treated cells, whereas TPC1-A1-N-treated cells displayed similar cholesterol content in CD63-positive compartments as the control.

**Fig. 5.**
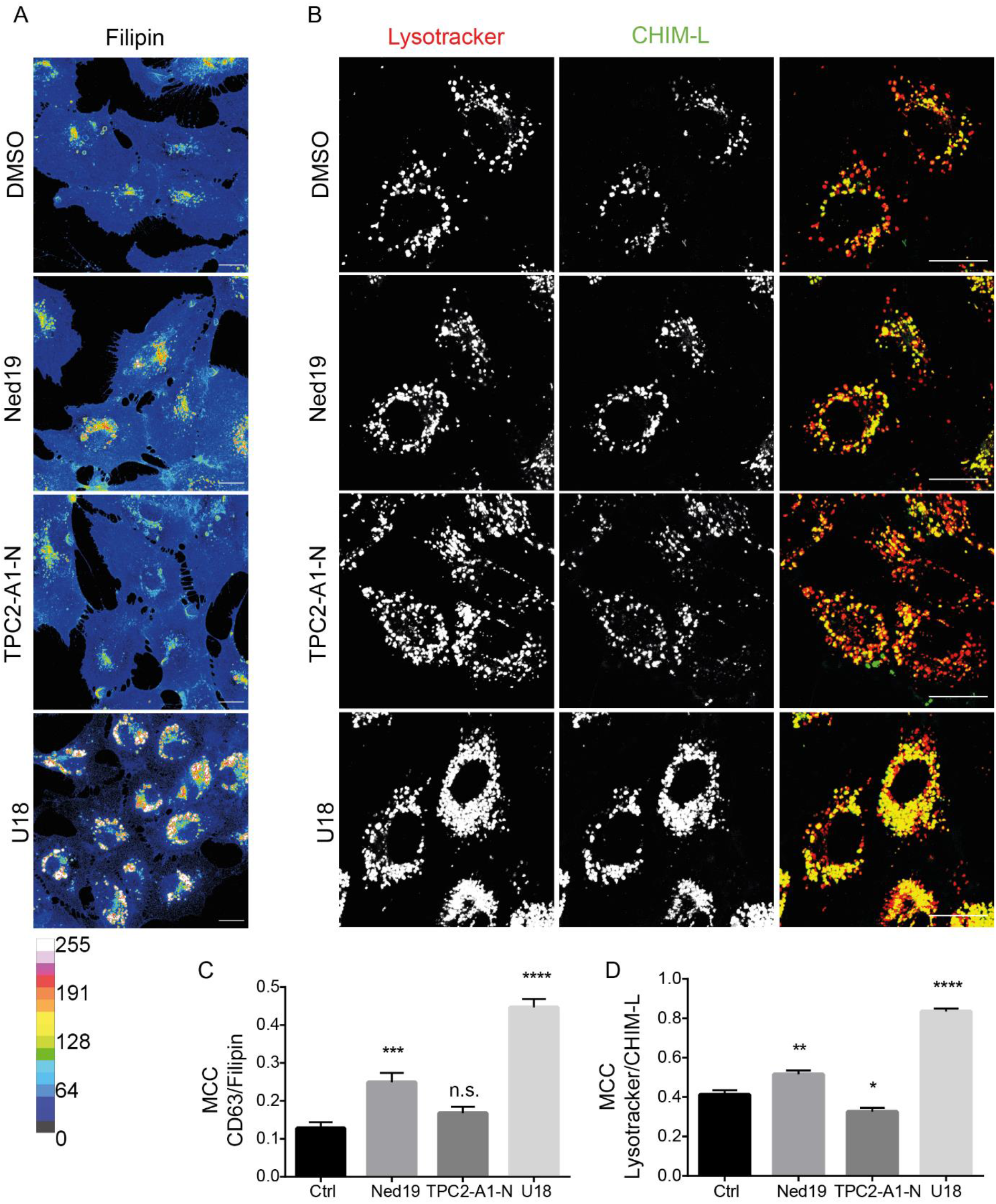
Targeting TPC2 affects endolysosomal cholesterol balance. Cells were either solvent-treated (Ctrl) or treated with the TPC-targeting compounds TPC1-A1-N (10 µM) and trans-Ned 19 (10 µM). U18666A (U18, 2µg/mL), the direct inhibitor of the endolysosomal cholesterol transporter NPC1, served as an internal reference. Filipin was used to detect cholesterol, CD63 or Lysotracker were used to identify endolysosomes. (A) Representative confocal images of treatment-induced endolysosomal cholesterol contents. Scale bars, 20 µm. To better visualize the endolysosomal cholesterol accumulation, filipin signal intensities were color-encoded (see calibration bar). (B) Representative confocal images of treatment-induced endolysosomal CHIM-L-AF488 contents. Scale bars, 20 µm. (C) The amount of endolysosomal cholesterol was assessed by calculating the Manders’ colocalization coefficients M1 of CD63 and filipin signals from z-stacks of individual cells. (D) The amount of endolysosomal CHIM-L-AF488 was assessed by calculating the Manders’ colocalization coefficients M1 of Lysotracker and CHIM-L-AF488 signals from z-stacks of individual cells. Data are presented as mean ± SEM of at least 41 cells from three independent experiments. Statistical analysis was performed by one-way ANOVA followed by Dunnett’s post-test (n.s., not significant, *p<0,05, **p<0.01, ***p<0.001, ****p<0.0001).

To study the effects of altered TPC2 functionality in living cells, we made use of the recently developed novel fluorescent cholesterol analog CHIM-L that mimics natural cholesterol behavior and has been successfully used to follow cholesterol dynamics *in vivo* (Matos *et al*, 2021). As shown in Fig. 5B, endocytosed CHIM-L-AF488 colocalized with the lysotracker signals in living cells. As expected, CHIM-L-AF488 accumulated when NPC1-mediated endolysosomal cholesterol egress was pharmacologically blocked with the NPC1 inhibitor U18. Importantly, endolysosomal CHIM-L-AF488 contents were elevated upon pre-treatment of cells with trans-Ned 19, whereas TPC2-A1-N treatment reduced the CHIM-L-AF488 content in LEL (Fig. 5D). Thus, both staining of endogenous cholesterol in fixed cells and with live cell analysis of the cholesterol analog CHIM-L-AF488 showed that TPC2 activity controls endolysosomal cholesterol egress.

Because efficient endolysosomal cholesterol egress depends on the low-pH-stabilized structural conformation of the cholesterol transporter NPC1 (Qian *et al*, 2020), we assessed whether our observation that TPC2 inhibition by trans-Ned 19 impairs endolysosomal cholesterol egress was caused by a concomitant endolysosomal alkalinization. We, therefore, determined the endolysosomal pH via ratiometric fluorescence microscopy (Kühnl *et al*, 2018). In line with previous observations (Gerndt *et al*, 2020), the pH values in cells treated with the TPC2 inhibitor trans-Ned 19 were lower than in the control cells, whereas TPC1-A1-N-mediated TPC2 activation caused a slight shift towards higher pH (Suppl. Fig. S3). We conclude that the changes in pH upon manipulation of TPC activity are likely not the cause of endolysosomal cholesterol accumulation.

### CD63 transport to WPB is governed by TPC2

In contrast to VWF and P-selectin, which are included already into nascent WPB, CD63 is sorted to mature WPB from the PM via endolysosomes. To explore the impact of TPC2 on CD63 downstream transport to WPB, we monitored this transport route via a well-established antibody uptake assay (Poeter *et al*, 2014). Therefore, cells were incubated with antibodies directed against an extracellular epitope of CD63. Subsequently, WPB were identified via VWF staining and the VWF-associated content of internalized anti-CD63 antibodies was quantified. As shown in Fig. 6A, anti-CD63 antibodies were reliably transferred onto WPB in control cells. In contrast, both siRNA-mediated ablation of TPC expression and trans-Ned 19-mediated TPC2 inhibition impaired significantly the PM-to-WPB transport of CD63, whereas TPC2 activation increased the CD63 antibody content in WPB. These results are in line with a specific role for TPC2 in maintaining endolysosome functionality (see also Fig. 2).

**Figure 6.**
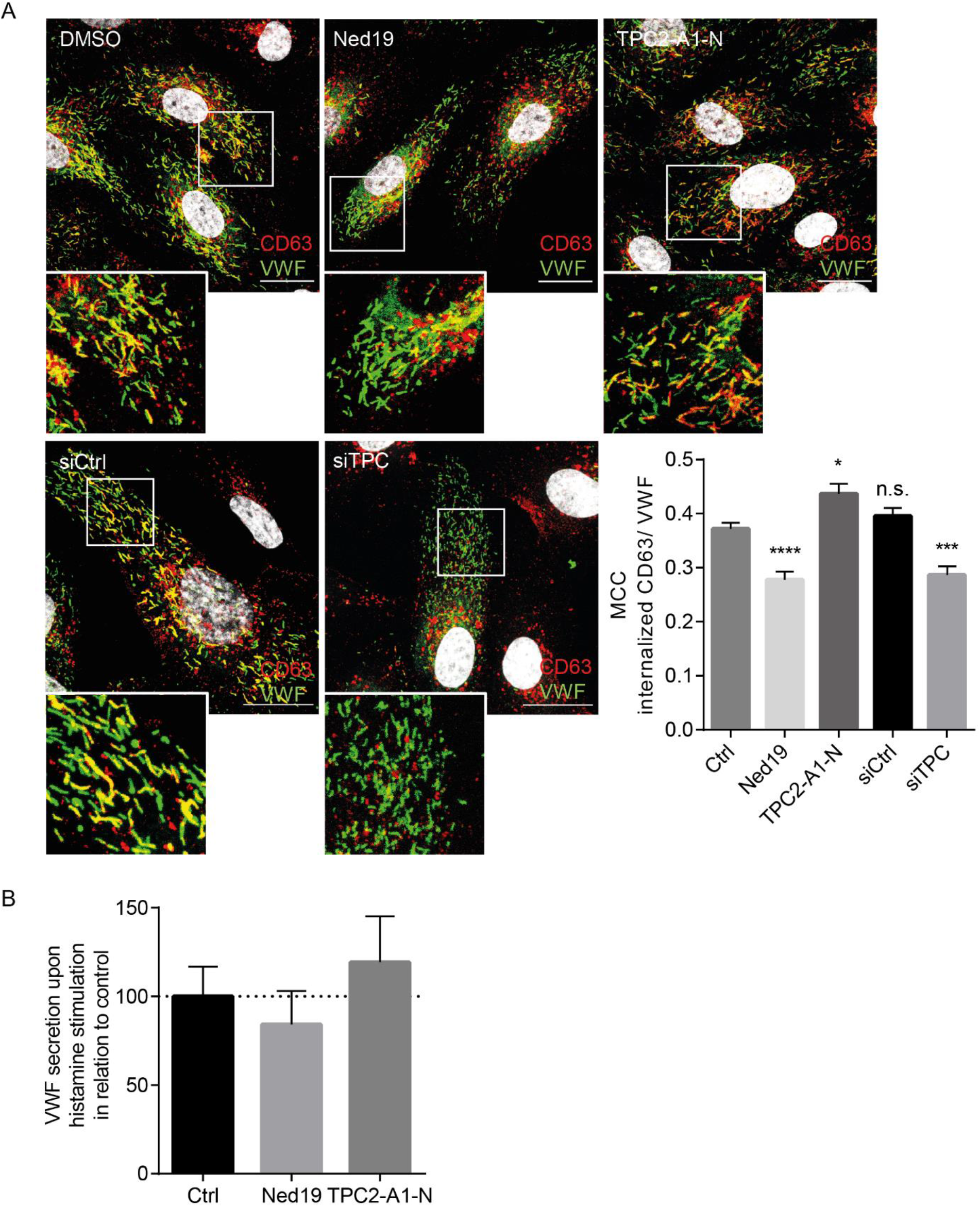
TPC2 is required for efficient CD63 transport to WPB but does not affect WPB exocytosis. (A) CD63 LEL to WPB transport was followed in cells treated with the TPC-targeting compounds (upper panel) or transfected with either control siRNA or TPC1/2 siRNA (lower panel). WPB were detected via VWF staining (green) and were analyzed for the amount of transferred anti-CD63 antibodies (red). Nuclei were visualized by DAPI (grey). Boxed areas indicate the regions magnified in the insets. Scale bars, 20 µm. To quantify the amount of anti-CD63 antibodies on WPB, Manders’ colocalization coefficients (MCC) were calculated from z-stack images of individual cells. Data represent mean ± SEM of at least 45 cells of three independent experiments and were analyzed by one-way ANOVA followed by Dunnett’s multiple comparison test (n.s., not significant, *p<0,05, **p<0.01, ***p<0.001, ****p<0.0001). (B) HUVEC were treated with the TPC2 inhibitor trans-Ned 19 or the activator TPC2-A1-N. WPB exocytosis was induced with histamine, and the amount of released VWF was quantified by ELISA. Note that TPC2 activity levels had no significant impact on VWF release.

We next investigated whether exocytosis of the WPB was controlled by TPC activity. Thus, we employed a well-established ELISA to quantify the release of VWF upon histamine stimulation as a read-out for pro-inflammatory, Ca^2+^-dependent WPB exocytosis. As shown in Fig. 6B, changing the activity levels of TPC2 was accompanied by only minor changes in the amount of released VWF which were not significant, arguing for a neglectable role of TPC2 in stimulated WPB exocytosis.

### TPC2 inhibition impairs histamine-evoked P-selectin cell surface presentation and leukocyte adhesion

To allow for immediate early leukocyte recruitment upon pro-inflammatory activation of the endothelium, the leukocyte receptor P-selectin and the stabilizing co-factor CD63 are both stored in WPB and are transferred to the cell surface via demand-driven WPB exocytosis. We, therefore, explored whether the inefficient loading of CD63 onto WPB impairs the inflammatory P-selectin/CD63 functions on the endothelial cell surface. The levels of P-selectin presented on the cell surface of trans-Ned 19-treated cells were significantly reduced (Fig. 7), suggesting that TPC2 activity controls the efficient delivery of CD63 to WPB which is in turn needed to stabilize P-selectin on the cell surface (Doyle *et al*, 2011; Poeter *et al*, 2014; Gerke, 2016). To further substantiate these findings, we explored whether leukocyte/endothelium interaction was affected by reducing TPC activity levels. Therefore, we treated HUVEC monolayers with trans-Ned 19 and determined the numbers of both rolling and adherent polymorphonuclear neutrophils (PMN) as well as their rolling velocity upon histamine activation under flow conditions. As shown in Fig. 8, histamine activation induced leukocyte rolling and adhesion. Importantly, TPC2 inhibition clearly impaired leukocyte recruitment. In all conditions, leukocyte/HUVEC interaction was abolished in the presence of anti-P-selectin antibodies, ruling out the involvement of other adhesion molecules, such as E-selectin.

**Fig. 7.**
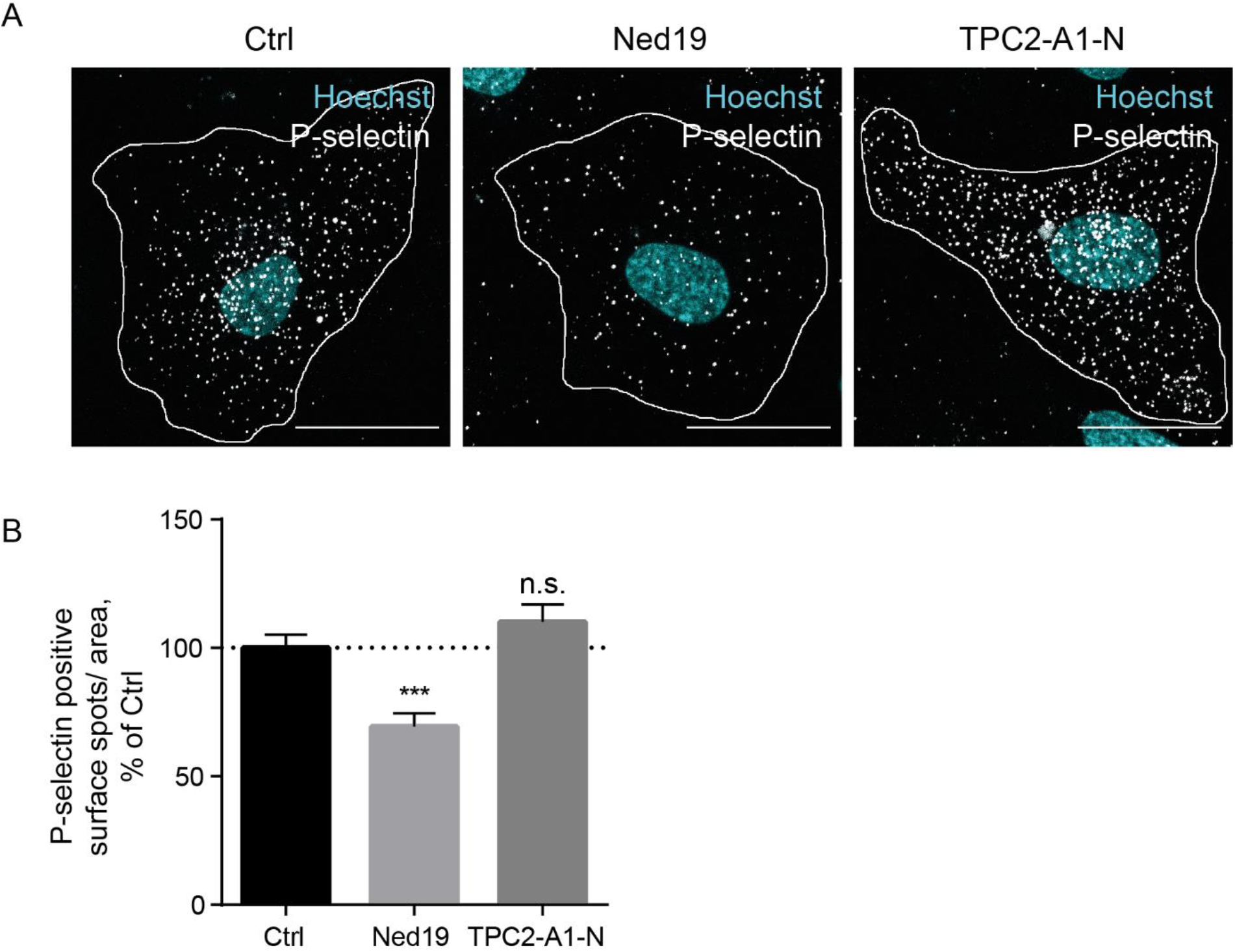
Targeting TPC2 activity affects histamine-evoked P-selectin cell surface presentation in HUVEC. (A) Representative images of cell-surface P-selectin pools of control cells and cells treated with either trans-Ned 19 (10 µM) or TPC2-A1-N (10 µM). Dashed lines outline cells of interest. Bars, 20 µm. (B) Quantitative analysis of P-selectin surface signals confirmed the altered cell surface levels of P-selectin. Data represent mean ± SEM of at least 22 cells per condition from three independent experiments and were analyzed by one-way ANOVA followed by Dunnett’s multiple comparison test (n.s., not significant, ***p<0.001).

**Figure 8.**
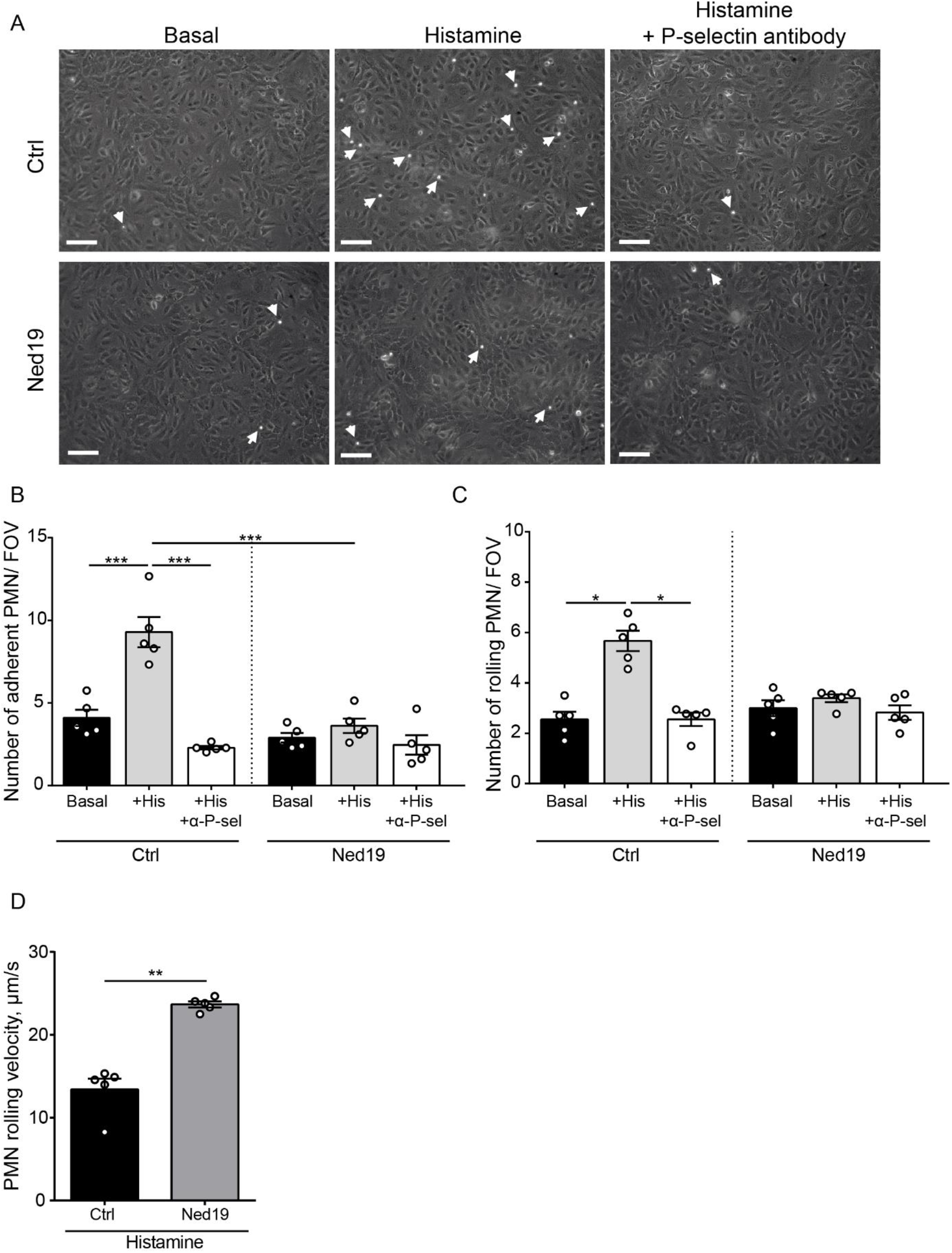
Endothelial TPC2 inhibition impairs histamine-evoked leukocyte recruitment. Adhesion of primary human neutrophils to unstimulated (basal), histamine-activated or anti-P-selectin-antibody-blocked HUVEC monolayers under flow. (A) Representative images after histamine treatment of control (top) and trans-Ned19-treated HUVEC monolayers (bottom). Arrows mark firmly attached leukocytes. Scale bars, 100 μm. Analysis of (B) PMN adhesion, (C) rolling, and (D) rolling velocities on control or trans-Ned 19-treated HUVEC monolayers. Data represent means ± SEM from five independent experiments (n=5). Data was statistically analyzed by one way ANOVA (adherent cells), Kruskal-Wallis test (rolling cells) and Mann-Whitney test (rolling velocity) (*p<0,05, **p<0.01, ***p<0.001).

## Discussion

Even though Ca^2+^ release from typical large Ca^2+^ stores like the endoplasmic reticulum is essential for signal transduction, smaller Ca^2+^ stores like LEL that are scattered throughout the cytosol seem to play a crucial role in local regulation events. Endolysosomal Ca^2+^ release affects, among others, membrane fusion, fission and local membrane trafficking (Yang *et al*, 2019; Morgan *et al*, 2011; Marchant & Patel, 2015; Xu & Ren, 2015). Therefore, we focused on the Two Pore channel (TPC) members TPC1 and TPC2. TPCs are endolysosomal non-selective cation channels that are involved in endolysosomal processes and cargo transport (Grimm *et al*, 2017). Interestingly, TPC ion selectivity has been revealed to depend on the activator: phosphatidylinositol 3,5-bisphosphate [PI(3,5)P2], an abundant endolysosomal phospholipid, elicits Na^+^-selective currents, whereas the second messenger nicotinic acid adenine dinucleotide phosphate (NAADP) elicits Ca^2+^ currents, leading to large increase in [Ca^2+^]i (Gerndt *et al*, 2020). Mobilization of endolysosomal Ca^2+^ is also induced by the recently developed synthetic TPC2 activator TPC2-A1-N (Gerndt *et al*, 2020), and NAADP binding and TPC2-mediated Ca^2+^ release is inhibited by the synthetic NAADP analog trans-Ned19 (Pitt *et al*, 2010; Naylor *et al*, 2009).

Our findings that the morphology of CD63 and LAMP2-positive endolysosomes was affected by the simultaneous knockdown of TPC1 and TPC2 confirmed a role for at least one of the channels in the maintenance of this compartment in HUVEC. Whereas TPC1 is predominantly localized on earlier endosomes, TPC2 is mainly found on endolysosomes, and both TPCs might sequentially function along the endolysosomal pathway (Morgan *et al*, 2011). Our observations that heterologously expressed TPC1 colocalized with markers of transferrin-containing endosomes but not with endolysosomal markers are well in agreement with these previous reports and argue against a role for TPC1 in the regulation of endolysosomes. A specific role for TPC2 to mediate endolysosome-related trafficking events was further supported by our observations that the LEL-localized TPC2-based Ca^2+^ sensors could be pharmacologically induced to transmit changes in Ca^2+^ levels upon addition of TPC2-A1-N (Gerndt *et al*, 2020). The absence of response of the pore-dead version of the TPC2-based Ca^2+^ sensor to TPC2-A1-N further confirmed that the Ca^2+^ fluxes were released via TPC2.

TPC2 dysregulation has been reported to cause a range of different pathological conditions and is observed in different diseases including Niemann-Pick disease type C1 (Lloyd-Evans *et al*, 2008), fatty-liver disease (Grimm *et al*, 2014) and neurodegenerative diseases such as Alzheimer’s and Parkinson’s (Feng & Yang, 2017). We observed that the endolysosomal cholesterol balance responded to a pharmacological manipulation of TPC2 function, with trans-Ned 19 treated HUVEC displaying cholesterol accumulation in CD63-positive compartments (Fig. 5). TPC1-A1-N-treated HUVEC displayed unchanged levels of cholesterol as indicated by filipin staining. Analysis of living cells exposed that endolysosomal levels of the novel cholesterol analog CHIM-L were affected by manipulated TPC2 activity as well. Trans-Ned 19 treated HUVEC showed CHIM-L accumulation in LEL, whereas TPC2-A1-N treatment even slightly reduced the amount of CHIM-L in LEL. Taken together, our data indicates that TPC2 activity is required for endolysosomal cholesterol egress, in line with what had been seen in the TPC2 KO mouse (Grimm *et al*, 2014).

Our findings that both siRNA-mediated ablation of TPC1/2 expression and pharmacological inhibition of TPC2 with trans-Ned 19 impaired the transfer of CD63 onto WPB significantly argue for a specific role for TPC2 and TPC2-mediated endolysosomal Ca^2+^ release in the delivery of cargo to WPB. Stimulated WPB release, on the other hand, was not measureably affected. This fits in with the fact that TPC were not detected on WPB, which is also in line with a currently published proteomic analysis of WPB that confirms the absence of WPB-localized TPC1 and TPC2 (Holthenrich *et al*, 2019). Interestingly, TPC2 is present and functional on another kind of lysosome-related organelles, the melanosomes (Ambrosio *et al*, 2016). The finding that the WPB do not contain TPC2, even though they share several endolysosomal features such as acidic pH, is most likely due to the fact that they bud from the Golgi and only mix to a limited extent with endolysosomes (Gerke, 2016; McCormack *et al*, 2017).

To ensure a tailored response toward inflammatory challenges, the endothelial cell surface is non-adhesive under resting conditions and only presents selectins upon pro-inflammatory induction. While expression of E-selectin requires de novo protein biosynthesis, the initial time gap is bridged via induced secretion of P-selectin from endothelial WPB (Subramaniam *et al*, 1993). Thus, WPB act as emergency kits that release their content to promote the immediate early inflammatory response until a more pronounced inflammatory activation of the endothelium is established (Schillemans *et al*, 2019).

Although both P-selectin and CD63 work together in leukocyte attraction and are present in WPB, they are sorted onto WPB at different stages of WPB genesis and only form complexes on the endothelial cell surface (Doyle *et al*, 2011; Harrison-Lavoie *et al*, 2006; Poeter *et al*, 2014). P-selectin is incorporated into nascent WPB that bud from the Golgi. The tetraspanin CD63, a marker protein for acidic endolysosomes, is only acquired by mature WPB from these compartments. While the importance of WPB formation and exocytosis for the proper inflammatory endothelial response is immediately evident, the delivery of cargo to WPB represents a further layer of regulation. P-selectin is stabilized on the endothelial cell surface via binding to the tetraspanin CD63 (Doyle *et al*, 2011). Indeed, we and others have shown that proper CD63 delivery from endolysosomes to WPB is a critical step in the establishment of the adhesive endothelial cell surface in vitro as well as in the murine model (Poeter *et al*, 2014; Kobayashi *et al*, 2000; Doyle *et al*, 2011). In this regard, it is interesting to note that our previous findings also revealed that impaired physiological functionality of late endosomes by pharmacologically inducing endolysosomal cholesterol build-up decreases the levels of CD63 delivery to WPB, causing a loss in P-selectin/CD63 cell surface presentation, and, in turn, severely reduced leukocyte recruitment (Poeter *et al*, 2014; Heitzig *et al*, 2018). Our results here revealed that TPC2 activity controlled the efficient delivery of CD63 to WPB and that inefficient loading of the P-selectin stabilizer CD63 onto WPB seen in cells treated with the TPC2 inhibitor trans-Ned 19 was associated with lower P-selectin levels on the endothelial cell surface and a concomitant impaired P-selectin-dependent leukocyte recruitment under flow. Next to the role of CD63 in the PM localization of P-selectin, CD63 also coordinate the vascular endothelial growth factor receptor 2 (VEGFR2)-β1 integrin complex formation and function at the cell surface (Tugues *et al*, 2013). Changes within the complex were shown to affect endothelial migration or invasion as well as tumor-associated angiogenesis (Heitzig *et al*, 2017).

The identification of regulatory elements is not only important for our understanding of basic endothelial cell functions but most likely also offers therapeutic points of attack. In this regard, juxta-endolysosomal Ca^2+^ concentration and its effect on endolysosomal trafficking towards WPB brings TPC into the focus as a potential target to treat several diseases including cancer (Heitzig *et al*, 2017) and chronical inflammation.

## Material and methods

### Antibodies and labeling reagents

The following primary antibodies were used: Mouse anti-CD63-TRITC (sc-5275, 1:100) and mouse anti-LAMP2 (sc-18822, 1:50) were obtained from Santa Cruz, rabbit anti-VWF (A0082, 1:1000) and rabbit anti-VWF-HRP (P0226, 1:8000) used for sandwich ELISA were purchased from Dako, sheep anti-VWF (ab11713, 1:200-400) and rabbit anti-TPC1 (ab94731, 1:200) were from Abcam, rabbit anti-TRPML1 was from Thermo Fisher Scientific (PA1-464741, 1:200), rabbit anti-TPC2 from Alomone Labs (ACC-072, 1:300), mouse anti-β-actin from Sigma (A5441, 1:5000) and sheep anti-P-selectin/CD62P from R+D Systems (AF137). The mouse anti-CD63 antibody H5C6c (1:300), developed by J.T. August and J.E.K. Hildreth, The Johns Hopkins University School of Medicine, was obtained from the Developmental Studies Hybridoma Bank, created by the NICHD of the NIH and maintained at The University of Iowa, Department of Biology, Iowa City, IA 52242.

For immunofluorescence analysis, the secondary antibodies were used at a dilution of 1:400. Donkey anti-mouse AF488 and goat anti-rabbit AF 488 were purchased from Invitrogen, donkey anti-mouse AF 647 was from Dianova, donkey anti-sheep AF 594 and AF 647 were obtained from Molecular Probes. DAPI and Filipin complex from Streptomyces filipinensis were purchased from Sigma.

For immunoblotting, the secondary antibodies donkey-α-mouse (H+L) and goat-α-rabbit (H+L) coupled to IRDye 680CW or IRDye 800CW (LI-COR) were used at a dilution of 1:5000.

### Compounds targeting and measuring Ca^2+^ release

Bafilomycin A1, ionomycin, and trans-Ned 19 were obtained from Cayman Chemicals. Oregon green 488-labeled dextran (OG, 10 kDa) was from Thermo Fisher Scientific, Flamma(R)-648-labeled dextran (Flamma, 10 kDa), Fluo-4-AM (Fluo4, ABD-20551), and FuraRed-AM (FuraRed, ABD-21048) were purchased from Biomol. The TPC2-selective agonist TPC2-A1-N (Gerndt *et al*, 2020) was provided by Franz Bracher, Munich.

### siRNA and plasmids

Gene-specific siRNA targeting TPC1 or TPC2 was from Dharmacon (L-010710-00-0005, L-006508-00-0005), non-targeting control siRNA was from Qiagen (1027281).

HsTPC1:mRFP, HsTPC2(L564P):mCherry (wild-type), TPC2(L564P):GCaMP6s (wild-type), and TPC2(L265P/L564P):CaMP6s (pore-dead) were provided by Christian Grimm (Munich). To generate wild-type TPC2(L564P):GCaMP6s and pore-dead TPC2(L265P/L564P):CaMP6s fused to mApple, TPC2(L564P):GCaMP6s and TPC2(L265P/L564P):CaMP6s sequences were amplified (forward primer, 5’-TCC<u>GAATTC</u>AATGGGTTCTCATCATCATCATCATCA-3’, reverse primer, 5’-GAT<u>ACCGGT</u>TGCAACTTCGCTGTCATCATTTGTACAAAC-3’) and subcloned into mApple N1-vector (Addgene #54567) via EcoRI und AgeI sites.

### Cell culture and transfection

Human umbilical vein endothelial cells (HUVEC) were obtained from Promocell and maintained as previously described (Heitzig *et al*, 2017). HUVEC were cultured in HUVEC medium composed of 50% Endothelial Cell Growth Medium 2 with Endothelial Cell Growth Medium 2 SupplementMix (C-39216, PromoCell) and 50% Medium 199 (Sigma) supplemented with 10% Fetal Bovine Serum Advanced (Capricorn Scientific), 30 µg/mL gentamycin (Sigma) and 15 ng/mL amphotericin B (Biochrom) at 5% CO_2_ and 37 °C on Corning CellBIND culture dishes. Coverslips and multiwell plates were coated with collagen type I (Advanced Biomatrix). Cells were transfected using the Amaxa HUVEC Nucleofector kit (Lonza) according to the manufacturer’s protocol (2-5 µg plasmid DNA, 400 pmol siRNA duplex per 3×10^6^ cells) and were analyzed 48 h post transfection.

### Quantitative real-time reverse transcription-PCR (qPCR)

Total cellular RNA was isolated with the RNeasy Kit (Qiagen) and was reverse-transcribed using the High-Capacity cDNA Reverse Transcription Kit (Thermo Fisher Scientific) according to the manufacturers’ instructions. qPCR was performed on a Roche LightCycler 480 using Brilliant III Ultra-Fast SYBR Green QPCR Master Mix (Agilent Technologies) and Qiagen Quantitect primers (Hs_ACTB_1_SG, QT00095431; Hs_GAPDH_2_SG, QT01192646; Hs_TPC1_1_SG, QT00088760; Hs_TPC2_1_SG, QT00084763). Relative gene expression levels were analyzed using the delta-delta Ct method (Livak & Schmittgen, 2001).

### Western blotting

Cells were lysed for 2 h in lysis buffer (62.5 mM Tris, pH 6.8, 10% glycerol, 5% β-mercaptoethanol, 4% sodium dodecyl sulfate, 0.002% bromophenol blue) on a shaker at 400 rpm at room temperature. Lysates were boiled for 10 min, proteins were separated on 12% gels via SDS-PAGE, and transferred to Amersham™ Protran® Western blotting nitrocellulose membranes (0.45 µm pore size, Sigma). Proteins were detected and quantified using the appropriate primary and secondary antibodies and an Odyssey infrared imaging system (LI-COR Biosciences).

### Confocal immunofluorescence analysis

HUVEC grown on collagen-coated coverslips were fixed for 10 min in 4% paraformaldehyde (PFA) at room temperature, permeabilized for 2 min with 0.2% Triton X-100 in PBS^++^, blocked for 15 min with 2% bovine serum albumin (BSA) in PBS^++^, and incubated for 1 h with the respective primary antibodies diluted in 2% BSA in PBS^++^, followed by incubation with appropriate secondary antibodies for 1 h. Nuclei were visualized with DAPI (1:1000 in PBS^++^, 10 min). Imaging of fixed cells and live-cell imaging was performed on an LSM 800 or LSM 780 microscope (Carl Zeiss, Jena, Germany) equipped with a Plan-Apochromat 63x/ 1.4 oil immersion objective. Confocal z-stack images and Image J software utilizing the JACoP plugin (Bolte & Cordelieres, 2006) for Fiji (Schindelin *et al*, 2012) was used to determine the Manders’ colocalization coefficients.

### CD63 antibody uptake

Uptake assays were performed as described previously (Poeter *et al*, 2014). In brief, cells cultured on collagen-coated coverslips were treated as indicated, incubated for 24 h in HUVEC medium containing anti-CD63 antibodies (H5C6c, 1:300), and fixed. Cells were imaged on an LSM800 Airyscan microscope (Carl Zeiss, Jena, Germany).

### Quantification of VWF secretion by sandwich ELISA

VWF levels in HUVEC lysates and culture media were measured as published previously (Disse *et al*, 2009; Chehab *et al*, 2017; Nguyen *et al*, 2020). In brief, cells grown on collagen-coated 24-well plates were incubated in basal assay medium (M199, 30 µg/mL gentamycin, 15 ng/mL amphotericin B, 2% BSA) for 20 min. Then the medium was collected and replaced with stimulation medium (basal assay medium containing 100 µM histamine, Sigma) for 20 min. Cells were then placed on ice and were lysed in lysis medium (basal assay medium, 1x protease inhibitor cocktail complete with EDTA (Roche), 0.1% Triton X-100) for 30 min.

VWF contents of basal and stimulation media, cell lysates, and VWF standard samples ranging from 0-1 ng VWF were measured in quadruplicates by a sandwich ELISA. Flat-bottom microtiter plates (Corning) coated with capture antibody solution (diluted 1:1000 in PBS) at 4°C for 16 h were washed with washing buffer (PBS, 0.5% Tween 20) and were subsequently blocked with 1.5% BSA in washing buffer at room temperature for 2 h. Samples were added and incubated at room temperature for 2 h. Plates were then washed with washing buffer and were incubated with HRP-labeled detection antibody solution (diluted 1:8000 in washing buffer with 1.5% BSA added) at room temperature for 2 h. Plates were washed and the substrate solution (TMB substrate Kit, Thermo Fisher Scientific # 34021) was added. After 20 min, 2 M sulfuric acid was added to stop the reaction. Absorbances were read at 450 nm in a CLARIOstar Plus plate reader (BMG Labtech). The average of the quadruplicates was used to calculate VWF concentrations in the media and the cell lysates.

### Endosomal pH measurement

Ratiometric fluorescence microscopy and calculation of endosomal pH were performed as previously published (Kühnl *et al*, 2018). Briefly, cells were pulsed with OG488-labeled and Flamma-labeled dextran in HUVEC medium at 37 °C for 1 h, followed by a 1 h chase. Cells were then washed and kept in 10 mM HEPES-buffered Hanks’ balanced salt solution (HBSS). Epifluorescence signals of z-stacks comprising 3-4 focal planes were acquired on a Zeiss LSM 780 microscope equipped with a Plan Apochromat 40x/1.4 Oil DIC M27 objective. Endosomal pH values were determined utilizing a calibration curve from HUVEC sequentially exposed for 10 min at 37°C to calibration solutions (20 mM HEPES, 143 mM KCl, 5 mM glucose, 1 mM MgCl_2_, 1 mM CaCl_2_, 10 µM nigericin) buffered to a pH ranging from 4.7 to 6.0.

### Endolysosomal calcium release measurement

HUVEC were washed with assay buffer (HBSS, 10 mM HEPES) and incubated with the Ca^2+^ indicators Fluo4 and FuraRed diluted to 0.5 µM each in assay buffer at 37°C for 25 min. Then, the indicator solution was replaced with assay buffer and the cells were transferred onto a Zeiss LSM 780 confocal microscope equipped with a 40x immersion oil objective. Cells were stimulated with 0.25 µM bafilomycin A1 or the TPC2 ligands as indicated. The maximum system response was determined by adding 0.5 µM ionomycin.

Epifluorescence signals were recorded every second and ratios of Fluo4/FuraRed fluorescence were calculated. Endosomal Ca^2+^ release was determined by subtracting baseline Fluo4/FuraRed ratios from the maximum Fluo4/FuraRed ratios after stimulation.

### Cholesterol visualization and colocalization analysis

To detect cholesterol, cells were fixed, blocked, and incubated with filipin (1.25 mg/mL in blocking buffer) for 2 h. Endolysosomal compartments were visualized with α-CD63 antibody (DSHB H5C6; 1:300 in blocking buffer). As an alternative to filipin staining for visualizing cholesterol distribution in HUVEC, we employed the clickable cholesterol analogue CHIM-L (Rakers *et al*, 2018; Matos *et al*, 2021). CHIM-L and the alkyne-fluorophore click-iT DIBO Alexa Fluor^®^ 488 (Thermo Fischer Scientific) were pre-clicked in a bio-orthogonal click reaction. Cells were incubated with 200 nM of the resulting CHIM-L:AF-488 overnight. To identify endolysosomes, lysoTracker Red DND-99 (Thermo Fisher, 75 nM) was added to the cells for 1 h at 37 °C. Live-cell imaging was performed in imaging buffer (FluoroBrite DMEM (Thermo Fisher), supplemented with 10 mM Hepes, 1 mM MgCl_2_, 0.9 mM CaCl_2_).

### Leukocyte adhesion assay

Leukocyte adhesion was assessed as published previously (Poeter *et al*, 2014). In brief, human neutrophils were isolated from human whole blood using Pancoll (pan-biotech) gradients according to the manufacturer’s protocol and as previously described (Rossaint *et al*, 2011). HUVEC grown to confluency on a 35 mm tissue culture dish (Corning) were incubated in media containing trans-Ned19 (10 µM) or the solvent control overnight. Monolayers were activated with 200 nM histamine for 20 min. To verify that leukocyte adhesion was dependent on P-selectin, monolayers were exposed to P-selectin antibodies for 30 min at 37 °C and capture was performed in the presence of 2 µg/mL anti-P-selectin antibody. PMN rolling and adhesion were determined using a parallel plate flow chamber assembly (Glycotech, Gaithersburg, MD, USA) perfused with isolated PMNs (2×10^5^ cells/mL) at a constant sheer stress of 0.55 dyne/cm^-2^ for 6 min (Schneider-Hohendorf *et al*, 2014). Movies were obtained with an inverted TS100 transmission light microscope (Nikon) equipped with a 10x/0.25 objective and a digital camera (Pixefly, Cooke Corporation, Romulus, MI, USA). Rolling and adherent cells were analyzed using ImageJ software. Rolling velocity (µm/s) was calculated by measuring the time required for a cell to roll across the distance covered in the optical field of view (FOV). The use of HUVEC and the isolation of human neutrophils from human whole blood was approved by the ethics committee of the University of Münster and the medical association of Westphalia-Lippe (ÄKWL).

### Statistical analysis

A priori power analysis (G*Power 3.1) was used to estimate the required sample sizes. A Gaussian distribution was tested through D’Agostino-Pearson omnibus K2 normality test.

Data were expressed as mean values ± standard error of the mean (SEM). Statistical significance of differences was evaluated by two-tailed unpaired Student’s *t*-test or one-way analysis of variance (ANOVA), followed by Dunnett’s multiple comparison test, using GraphPad Prism version 6.0 (GraphPad software, San Diego, CA, USA). P < 0.05 was considered significant.

## Author Contributions

Conceptualization and methodology, U.R., N.H.; validation, formal analysis, investigation, data curation, J.G., N.H., K.T., J.N., A.L.L.M.; resources, E.K.K. C.G., U.R., V.G., J.R, T.W., F.G., F.B.; writing—original draft preparation, N.H., J.G; writing—review and editing, C.G., N.H., V.G., J.G., S.S., U.R.; visualization, J.G., N.H.; supervision, N.H., U.R., C.G.; project administration, U.R.; funding acquisition, U.R, S.S., F.G., F.B., C.G.. All authors have read and agreed to the published version of the manuscript.

## Acknowledgments

We thank Katharina Ludewig for help with the CD63 antibody transport assays.

## The paper explained

### Problem

Inflammatory diseases are characterized by excessive neutrophil activation. Because interfering with the adherence of leukocytes to the endothelial cells might restore a balanced neutrophil recruitment, a mechanistic understanding of neutrophil-endothelial cell interactions is required to identify points of intervention. Upon inflammatory activation, endothelial cells present adhesion molecules on their cell surface to attract leukocytes. Induction by histamine leads to a rapid transport of the leukocyte adhesion receptor P-selectin and the stabilizing cofactor CD63 from specialized intracellular storage organelles, the Weibel-Palade bodies (WPB), to the plasma membrane.

### Results

We report that the endolysosomal two pore channel 2 (TPC2) promotes the proper loading of CD63 onto WPB in primary human endothelial cells. Interfering with TPC2 functionality via siRNA-mediated TPC2 knockdown or pharmacological inhibition of TPC2 reduced the amount of WPB-associated CD63 which was instead retained in endolysosomes. Pharmacological inhibition of TPC2 impaired P-selectin presentation on the surface of histamine-stimulated endothelial cells and severely reduced leukocyte recruitment under flow.

### Impact

Our findings establish the relevance of CD63 transport from endolysosomes onto WPB in the endothelial pro-inflammatory conversion and identify the endolysosomal TPC2 ion channel as a novel regulatory element and a potential point of attack to target the endothelial interaction with leukocytes.

## References

Ambrosio AL, Boyle JA, Aradi AE, Christian KA & Di Pietro SM (2016) TPC2 controls pigmentation by regulating melanosome pH and size. Proc Natl Acad Sci U S A 113: 5622–5627

Bolte S & Cordelieres FP (2006) A guided tour into subcellular colocalisation analysis in light microscopy. J Microsc 224: 13–232

Chehab T, Santos NC, Holthenrich A, Koerdt SN, Disse J, Schuberth C, Nazmi AR, Neeft M, Koch H, Man KNM, et al (2017) A novel Munc13-4/S100A10/annexin A2 complex promotes Weibel-Palade body exocytosis in endothelial cells. Mol Biol Cell 28: 1688–1700

Davis LC, Morgan AJ, Chen J-L, Snead CM, Bloor-Young D, Shenderov E, Stanton-Humphreys MN, Conway SJ, Churchill GC, Parrington J, et al (2012) NAADP Activates Two-Pore Channels on T Cell Cytolytic Granules to Stimulate Exocytosis and Killing. Curr Biol 22: 2331–2337

Disse J, Vitale N, Bader M-F & Gerke V (2009) Phospholipase D1 is specifically required for regulated secretion of von Willebrand factor from endothelial cells. Blood 113: 973–980

Doyle EL, Ridger V, Ferraro F, Turmaine M, Saftig P & Cutler DF (2011) CD63 is an essential cofactor to leukocyte recruitment by endothelial P-selectin. Blood 118: 4265–4273

Feng X & Yang J (2017) Lysosomal Calcium in Neurodegeneration. Messenger 5: 56–66

Gerke V (2016) Annexins A2 and A8 in endothelial cell exocytosis and the control of vascular homeostasis. Biol Chem 397: 995–1003

Gerndt S, Chen CC, Chao YK, Yuan Y, Burgstaller S, Rosato AS, Krogsaeter E, Urban N, Jacob K, Nguyen ONP, et al (2020) Agonist-mediated switching of ion selectivity in TPC2 differentially promotes lysosomal function. Elife 9: 1–63

Gimpl G & Gehrig-Burger K (2007) Cholesterol Reporter Molecules. Biosci Rep 27: 335–358

Grimm C, Chen CC, Wahl-Schott C & Biel M (2017) Two-pore channels: Catalyzers of endolysosomal transport and function. Front Pharmacol 8: 6–11

Grimm C, Holdt LM, Chen C-C, Hassan S, Müller C, Jörs S, Cuny H, Kissing S, Schröder B, Butz E, et al (2014) High susceptibility to fatty liver disease in two-pore channel 2-deficient mice. Nat Commun 5: 4699

Harrison-Lavoie KJ, Michaux G, Hewlett L, Kaur J, Hannah MJ, Lui-Roberts WWY, Norman KE & Cutler DF (2006) P-selectin and CD63 use different mechanisms for delivery to Weibel-Palade bodies. Traffic 7: 647–662

Heitzig N, Brinkmann BF, Koerdt SN, Rosso G, Shahin V & Rescher U (2017) Annexin A8 promotes VEGF-A driven endothelial cell sprouting. Cell Adh Migr 11: 275–287

Heitzig N, Kühnl A, Grill D, Ludewig K, Schloer S, Galla HJ, Grewal T, Gerke V & Rescher U (2018) Cooperative binding promotes demand-driven recruitment of AnxA8 to cholesterol-containing membranes. Biochim Biophys Acta - Mol Cell Biol Lipids 1863: 349–358

Holthenrich A, Drexler HCA, Chehab T, Naß J & Gerke V (2019) Proximity proteomics of endothelial Weibel-Palade bodies identifies novel regulator of von Willebrand factor secretion. Blood 134: 979–982

Jahidin AH, Stewart TA, Thompson EW, Roberts-Thomson SJ & Monteith GR (2016) Differential effects of two-pore channel protein 1 and 2 silencing in MDA-MB-468 breast cancer cells. Biochem Biophys Res Commun 477: 731–736

Kobayashi T, Vischer UM, Rosnoblet C, Lebrand C, Lindsay M, Parton RG, Kruithof EK & Gruenberg J (2000) The tetraspanin CD63/lamp3 cycles between endocytic and secretory compartments in human endothelial cells. Mol Biol Cell 11: 1829–1843

Kühnl A, Musiol A, Heitzig N, Johnson DE, Ehrhardt C, Grewal T, Gerke V, Ludwig S & Rescher U (2018) Late endosomal/lysosomal cholesterol accumulation is a host cell-protective mechanism inhibiting endosomal escape of influenza A virus. MBio 9: e01345–18

Livak KJ & Schmittgen TD (2001) Analysis of relative gene expression data using real-time quantitative PCR and the 2-ΔΔCT method. Methods 25: 402–408

Lloyd-Evans E, Morgan AJ, He X, Smith DA, Elliot-Smith E, Sillence DJ, Churchill GC, Schuchman EH, Galione A & Platt FM (2008) Niemann-Pick disease type C1 is a sphingosine storage disease that causes deregulation of lysosomal calcium. Nat Med 14: 1247–1255

Marchant JS & Patel S (2015) Two-pore channels at the intersection of endolysosomal membrane traffic. Biochem Soc Trans 43: 434–441

Matos ALL, Keller F, Wegner T, del Castillo CEC, Grill D, Kudruk S, Spang A, Glorius F, Heuer A & Gerke V (2021) CHIMs are versatile cholesterol analogs mimicking and visualizing cholesterol behavior in lipid bilayers and cells. Commun Biol 4

McCormack JJ, Lopes da Silva M, Ferraro F, Patella F & Cutler DF (2017) Weibel™Palade bodies at a glance. J Cell Sci 130: 3611–3617

McEver RP (2015) Selectins: Initiators of leucocyte adhesion and signalling at the vascular wall. Cardiovasc Res 107: 331–339 doi:10.1093/cvr/cvv154

Morgan AJ & Galione A (2021) Lysosomal agents inhibit store-operated Ca2+ entry. J Cell Sci 134

Morgan AJ, Platt FM, Lloyd-Evans E & Galione A (2011) Molecular mechanisms of endolysosomal Ca2+ signalling in health and disease. Biochem J 439: 349–378

Morgan AJ, Yuan Y, Patel S & Galione A (2020) Does lysosomal rupture evoke Ca2+ release? A question of pores and stores. Cell Calcium 86: 102139

Naylor E, Arredouani A, Vasudevan SR, Lewis AM, Parkesh R, Mizote A, Rosen D, Thomas JM, Izumi M, Ganesan A, et al (2009) Identification of a chemical probe for NAADP by virtual screening. Nat Chem Biol 2009 54 5: 220–226

Nguyen TTN, Koerdt SN & Gerke V (2020) Plasma membrane phosphatidylinositol (4,5)-bisphosphate promotes Weibel–Palade body exocytosis. Life Sci Alliance 3: e202000788

Phillipson M & Kubes P (2011) The neutrophil in vascular inflammation. Nat Med 17: 1381–1390 doi:10.1038/nm.2514

Pitt SJ, Funnell TM, Sitsapesan M, Venturi E, Rietdorf K, Ruas M, Ganesan A, Gosain R, Churchill GC, Zhu MX, et al (2010) TPC2 Is a Novel NAADP-sensitive Ca2+ Release Channel, Operating as a Dual Sensor of Luminal pH and Ca2+. J Biol Chem 285: 35039–35046

Poeter M, Brandherm I, Rossaint J, Rosso GG, Shahin V, Skryabin B V., Zarbock A, Gerke V & Rescher U (2014) Annexin A8 controls leukocyte recruitment to activated endothelial cells via cell surface delivery of CD63. Nat Commun 5: 3738

Qian H, Wu X, Du X, Yao X, Zhao X, Lee J, Yang H & Yan N (2020) Structural Basis of Low-pH-Dependent Lysosomal Cholesterol Egress by NPC1 and NPC2. Cell 182: 98-111.e18

Rakers L, Grill D, Matos ALL, Wulff S, Wang D, Börgel J, Körsgen M, Arlinghaus HF, Galla H-J, Gerke V, et al (2018) Addressable Cholesterol Analogs for Live Imaging of Cellular Membranes. Cell Chem Biol 25: 952-961.e12

Rossaint J, Spelten O, Kässens N, Mueller H, Van Aken HK, Singbartl K & Zarbock A (2011) Acute loss of renal function attenuates slow leukocyte rolling and transmigration by interfering with intracellular signaling. Kidney Int 80: 493–503

Rossaint J & Zarbock A (2013) Tissue-specific neutrophil recruitment into the lung, liver, and kidney. J Innate Immun 5: 348–357 doi:10.1159/000345943

Schillemans M, Karampini E, Kat M & Bierings R (2019) Exocytosis of Weibel–Palade bodies: how to unpack a vascular emergency kit. J Thromb Haemost 17: 6–18

Schindelin J, Arganda-carreras I, Frise E, Kaynig V, Longair M, Pietzsch T, Preibisch S, Rueden C, Saalfeld S, Schmid B, et al (2012) Fiji : an open-source platform for biological-image analysis. Nat Methods 9: 676–682

Schneider-Hohendorf T, Rossaint J, Mohan H, Böning D, Breuer J, Kuhlmann T, Gross CC, Flanagan K, Sorokin L, Vestweber D, et al (2014) VLA-4 blockade promotes differential routes into human CNS involving PSGL-1 rolling of T cells and MCAM-adhesion of TH17 cells. J Exp Med 211: 1833–1846

Subramaniam M, Koedam JA & Wagner DD (1993) Divergent fates of P-and E-selectins after their expression on the plasma membrane. Mol Biol Cell 4: 791–801

Tugues S, Honjo S, König C, Padhan N, Kroon J, Gualandi L, Li X, Barkefors I, Thijssen VL, Griffioen AW, et al (2013) Tetraspanin CD63 Promotes Vascular Endothelial Growth Factor Receptor 2-β1 Integrin Complex Formation, Thereby Regulating Activation and Downstream Signaling in Endothelial Cells in Vitro and in Vivo. J Biol Chem 288: 19060–19071

Xu H & Ren D (2015) Lysosomal Physiology. Annu Rev Physiol 77: 57–80

Yang J, Zhao Z, Gu M, Feng X & Xu H (2019) Release and uptake mechanisms of vesicular Ca 2+ stores. Protein Cell 10: 8–19 doi:10.1007/s13238-018-0523-x

Zhu MX, Ma J, Parrington J, Calcraft PJ, Galione A & Evans AM (2010) Calcium signaling via two-pore channels: Local or global, that is the question. Am J Physiol -Cell Physiol 298: 430–441 doi:10.1152/ajpcell.00475.2009

