## Supplemental Figures for "Leukocyte adhesion is governed by endolysosomal two pore channel 2 (TPC2)"

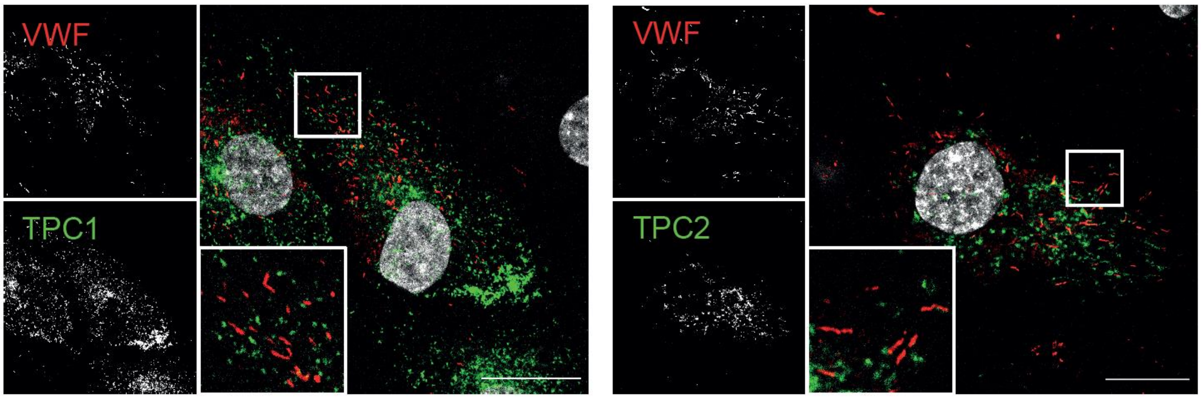


**Supplemental figure S1: TPC1 and TPC2 do not reside on Weibel-Palade bodies (WPB).** HUVEC transfected with TPC1-RFP (green, left) and TPC2-mCherry (green, right) were fixed and stained with an antibody against von-Willebrand Factor (VWF, red) and Dapi (grey). Even though WPB are in close proximity to endosomal TPC-positive compartments, the mature WPB do not colocalize with either TPC1 or TPC2. Images were taken with an LSM 800 airyscan microscope equipped with a 63x oil-objective (Zeiss). A representative single plane of a Z-stack image is depicted here. Scale bar: 20 µm.

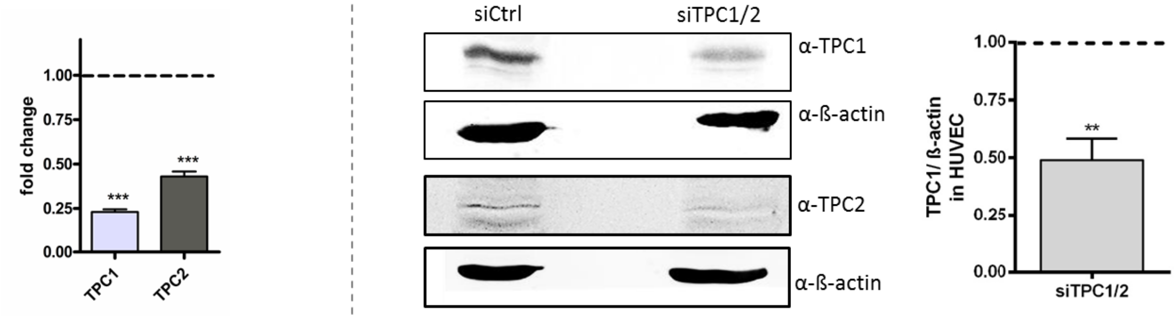


**Supplemental figure S2: TPC knockdown verification via qPCR (left) and western blot (right).** HUVEC were transfected with non-targeting siRNA (siCtrl) or TPC1/2 siRNA to reduce cation channel expression 48h post transfection. GAPDH and β-actin were used as housekeeping genes. Knockdown efficiency was statistically evaluated by Student's t-test (**p<0.01, ***p<0.001).

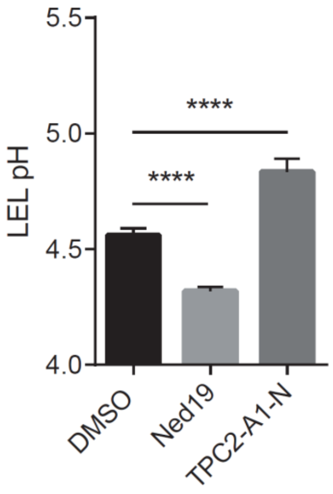


**Supplemental figure S3: Ratiometric in vivo pH measurements showing that endolysosomes were more acidified upon trans-Ned 19 inhibition of TPC2, whereas TPC2-A1-N-mediated TPC2 activation led to endolysosomal alkalinization.** Data are presented as mean ± SEM of 126 cells per condition from at least three independent experiments and were analyzed by one-way ANOVA followed by Dunnett's post-test (****$p<0.0001$).
